## Supplementary Material for "Controllability and cause in human collaboration"

### Supplementary Materials

#### Supplementary methods and results

##### Participants and recruitment

**MRI sample.** The analysed MRI data set originates from a larger sample with participants on a range of depression symptoms (findings to be reported elsewhere). To be included in the MRI study, participants were right-handed, aged 18-45 years, with normal or corrected to normal vision, and fluent in English. Participants were excluded if they had any contraindications for MRI scanning, current alcohol or drug dependency, current or past history of bipolar or psychosis disorder, or current use of psychoactive medication or recreational drugs. Interested individuals filled in a pre-screening online form hosted on Qualtrics. This online form included questions on demographics, MRI safety and a short depression questionnaire (Quick Inventory of Depressive Symptomatology, QIDS-SR) [1]. Participants were then invited to a pre-screening video call during which MRI safety was checked again, and during which part of the Structured Clinical Interview for DSM-5 Disorders was administered [2]. Participants were then invited to a single study visit. In this study, only participants with none or mild depressive symptoms (QIDS-SR total score < 11) were included.

**Online sample.** The study was advertised on the online platform Prolific (www.prolific.com). Participants were included into the study if they were aged 18-40 years old, had English as their first language, normal or corrected to normal vision, and at least 10 previous study submissions. Online participants could not move onto the main task if they failed the "system check" for the games. During this system check, participants practised the games and were instructed to do their best. For each of the games, participants had to produce five consecutive trials with sufficiently good game performance (defined as lower than 90% of largest absolute objective error from pilot data) to confirm that the games work correctly on their systems and that they paid attention. For each game, during task practice, participants had a maximum of 30 trials during which to produce those five good trials. Additionally, participants were excluded if after three attempts, they did not pass the multiple-choice test which checked their task understanding. After the first and second false attempt at this comprehension check, they were shown which questions they answered incorrectly and had to reread the instructions.

##### Experimental design

**Post-task debrief questionnaires**. After finishing the task, participants filled in a debrief questionnaire. One of the questions they answered was "In the study, how many times per game did you do badly on purpose?", which was answered on a Likert scale from "never", "less than once per game", 1, 2, …, 10, "More than 10 times per game". 5 acquired MRI participants (one excluded from analysis) did not fill in the debrief questionnaire because it was only added at a later stage. Online participants filled in a more extensive debrief questionnaire. One of the additional questions filled in only by the online sample was "If you did badly on purpose, how did it help you do each rating?", which for each rating (self, other, control) could be answered on a Likert scale from 1 – Not at all, 2, 3, 4, 5, 6 – Very much.

**Mood ratings and psychiatric questionnaires.** During the task, participants repeatedly rated their current mood. MRI participants also filled in a series of psychiatric questionnaires after the task and debrief questionnaire. Data from the mood ratings and psychiatric questionnaires were not analysed here and findings will be reported elsewhere.

**Mapping of self-performance.** On every trial, participants' objective game performance was transformed into a self-performance, according to the following logistic function:

$$Self(t) = 100 * \left(1 - \left(1 + e^{Slope(t) * (Shift(t) - Error(t))}\right)^{-1}\right) \quad \text{(S1)}$$

with

$$Slope(t) = \frac{\ln((1 - 0.55)^{-1} - 1)}{Error_{mean} - Error_{min}} \quad \text{(S2)}$$

and

$$Shift(t) = Error_{mean} + \ln\left(\left(1 - \frac{Self_{level}}{100}\right)^{-1} - 1\right) * Slope(t)^{-1} \quad \text{(S3)}$$

where *Self(t)* is the transformed self-performance (point score between 0 and 100) on a given trial *t*. *Error(t)* is the absolute objective performance error on trial *t*. In the rockslide game, for example, this was the absolute Euclidian distance (in pixels) between the target cross and the ball when participants pressed the button. If the participant pressed the button much too early or late, when the ball was not on the same slide as the target cross, their objective performance was always transformed to a *Self*=0 (active disambiguation trial, see supplementary Figure S4). The parameter *Slope(t)* determines the steepness of the mapping curve and reflects how sensitive the self performance is to the objective errors. It is determined based on the mean ($Error_{mean}$) and smallest error ($Error_{min}$) from the previous five trials, excluding AD trials. On the first five trials of a game block, this was based on the pre-task training trials of the respective game. This staircasing procedure was introduced because some participants' errors drifted over time. Calibrating the slope to the participant's previous error range ensured that the mapped self-performance has a constant variance across trials and is the same across participants. The *Shift(t)* parameter computes what error value is mapped onto the $Self_{level}$. $Self_{level}$ is the self-performance level pre-determined by our schedules (see section of schedules), and is the mean self-performance in the current task block. This is also the value that participants need to infer for their own self performance. The shift parameter depends on participants' mean errors across the previous five trials so that the $Self_{level}$ is given for participants' individual mean error. Calibrating the slope and shift parameters to the participants' recent error history ensured that the mapped self-performance remained constant and centred around the $Self_{level}$. Supplementary Figure S2 illustrates the logistic function as well as the staircasing for an example participant.

#### Behavioural analyses and results

**Accuracies of individual ratings and estimated feedback.** If participants are presented with very good feedback, they should believe that either they themselves or the other player did very well (or even both). However, there is an infinite number of possible combinations of self, other and control that can all lead to the same feedback. For example, a low feedback could result from one's own performance being low and a high control for oneself, or from the other's performance being low and the other having a high level of control. This means that while participants' individual ratings (or beliefs) might not be accurate, their ratings in combination should closely match the ambiguous feedback.

**Inferring self and other from active disambiguation.** Just as control can be inferred in the Control-Other phase, self-performance levels ("self") can be inferred from those trials during the Self-Other phase (see Supplementary Figure S6A left). Here, self can be inferred from the difference between normal and AD trials, taking into account the known control level. We tested whether participants used this logic (equation S4) with a regression model (equation S5):

$$Self = \frac{feedback\ difference}{Control} \tag{S4}$$

$$\begin{gathered} self \sim feedback\ difference + inverse\ control + \\ feedback\ difference * inverse\ control + previous\ self + \\ (feedback\ difference + inverse\ control + \\ feedback\ difference * inverse\ control + previous\ self|ID) \end{gathered} \tag{S5}$$

where *self* is the next trial's self rating. *Feedback difference* is the difference between normal and AD feedback, as observed on the current and previous trials. *Inverse control* is ln((current trial's control level)$^{-1}$), and *previous self* is the current trial's self rating. The hierarchical regression model was fit separately to participants' ratings, and to the respective control estimates of the active, ignorant and passive Bayesian learning models. Similar to the equivalent control regressions, this regression was fit to switch trials only as this is when the feedback difference can be observed. We fit this regression to participants' self ratings as well as the self estimates of the active, passive and ignorant Bayesian learning models. Results of this regression analysis are shown in Supplementary Figure S6B.

From the feedback equations it follows that participants could infer the other from the AD feedback accounting for the other's control. This is because on AD trials, the feedback shown is only based on the other's performance, weighted by the other's control. We tested whether participants used this logic (equation S6) using regressions fit to their other ratings (equation S7):

$$Other = \frac{AD\ feedback}{Other\ control} \tag{S6}$$

$$\begin{gathered} other \sim AD\ feedback + inverse\ other\ control + \\ AD\ feedback * inverse\ other\ control + previous\ other + \\ (AD\ feedback + inverse\ other\ control + \\ AD\ feedback * inverse\ other\ control + previous\ other|ID) \end{gathered} \tag{S7}$$

where other is the next trial's other rating. *AD feedback* is the feedback observed on the current AD trial. *Inverse other control* is the inverse of the control that the other player exerts, i.e. ln((1-Control)$^{-1}$). This regression was fit only to AD trials as this is when the AD feedback can be observed, and fit separately to the Self-Other and the Control-Other phase. In the Self-Other phase, *inverse other control* is computed with the known control level, while it is computed based on the control ratings in the Control-Other phase. *Previous other* is the current trial's other rating, i.e. the rating just before observing the AD feedback. This regression was fit separately to participants' other ratings, and the other estimates of the active, passive and ignorant learners. For the learning models, *inverse other control* was based on the control estimates from the respective model.

**Control effect on self and other ratings.** In the regression results shown in Figure 2E, we also find that overall, participants update their beliefs about the other (but not themselves) depending on the control level (main effect *1-Control* on other rating update: $F(1,65)=16.02$, $p<0.001$, $\eta^2=0.20$; *Control* on self-rating update: $F(1,65)=0.88$, $p=0.35$, $\eta^2=0.01$). The active learner showed a main effect of control on both self and other (*Control* on self, $F(1,65)=69.38$, $p<0.001$, $\eta^2=0.51$; *1-Control* on other, $F(1,65)=68.33$, $p<0.001$, $\eta^2=0.51$). Since we did not have hypotheses about this effect of control and the overall effect size is very small, we will not interpret it further.

#### Supplementary figures

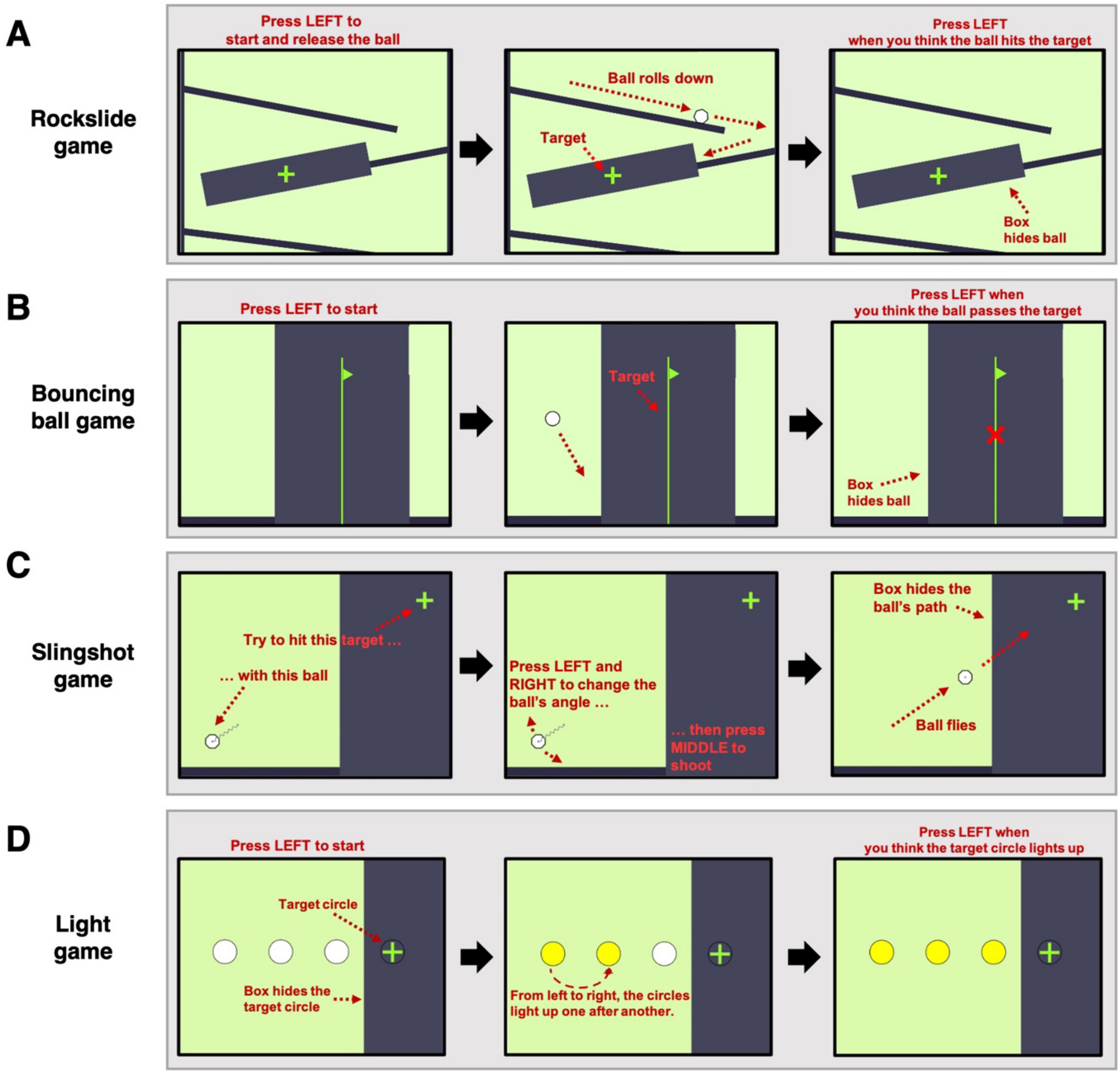


**Figure S1 (relating to Figure 1).** Games in the credit assignment task. **A)** In the Rockslide game, participants had to press the left button to release a ball that then rolled down a set of slides from the top. Participants then pressed the left button again to indicate when they thought that the ball passed the target cross. A black box hid the ball's path so that they could not see when the ball passes by the target exactly. The box was centred around the target cross. From trial to trial, the target cross changed its position slightly on the second slide. **B)** In the Bouncing ball game, participants pressed the left button to release a ball that would appear from the left and bounce towards the right. Participants pressed the left button again when they thought the ball passed the target flag. The black box hid the ball's path so that participants again were not able to tell when exactly the ball passed the target flag. On every trial, the target flag had a different horizontal position and the black box was centred around the target flag. **C)** In the Slingshot game, participants adjusted the angle of a ball (with left and right buttons) to hit a target cross. They pressed their middle button to shoot the ball. Again, a black box hid the ball's path so that participants were not able to see how accurately they shot the ball. On every trial, the target cross changed position vertically. **D)** In the Light game, the participants saw a set of white circles. When participants pressed the left button to start, the white circles lit up one after another from the left to the right. The fourth circle, the target circle, was hidden behind a black box so that participants did not see when it lit up. They had to press the button when they thought that the target circle would light up. On every trial, the latency between start (when participants pressed the left button the first time) and when the target circle would light up changed.

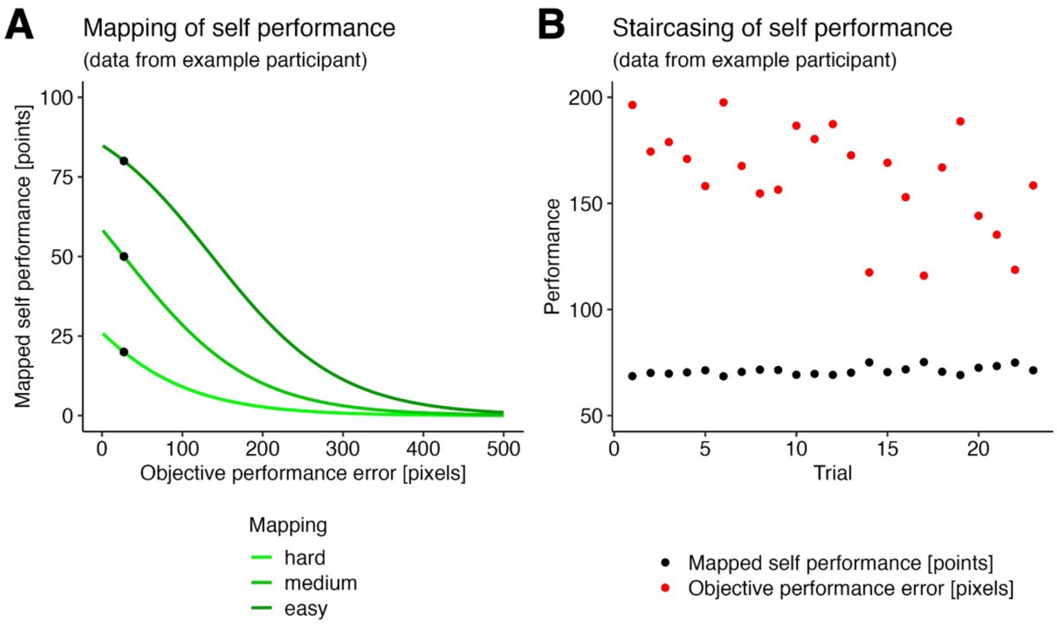


**Figure S2 (relating to Figure 1).** Transformation of objective game performance into self-performance. **A)** Mapping of objective performance error onto the self-performance. We used a logistic function to transform participants' objective performance errors into the self-performance. A lower objective performance error (i.e. higher accuracy) resulted in higher self-performance. Note that under different mapping conditions, the same objective performance error resulted in differently mapped self-performances (black dots). Values of an example participants are shown here. **B)** Staircasing of self-performance. Over time, this example participant drifted in their objective performance (red dots) and got better over time. By using a staircasing procedure, we transformed their drifting performance into a stable self-performance (black dots). This meant that even if participants objectively got better or worse at the games over time, the mapped self-performance remained the same.

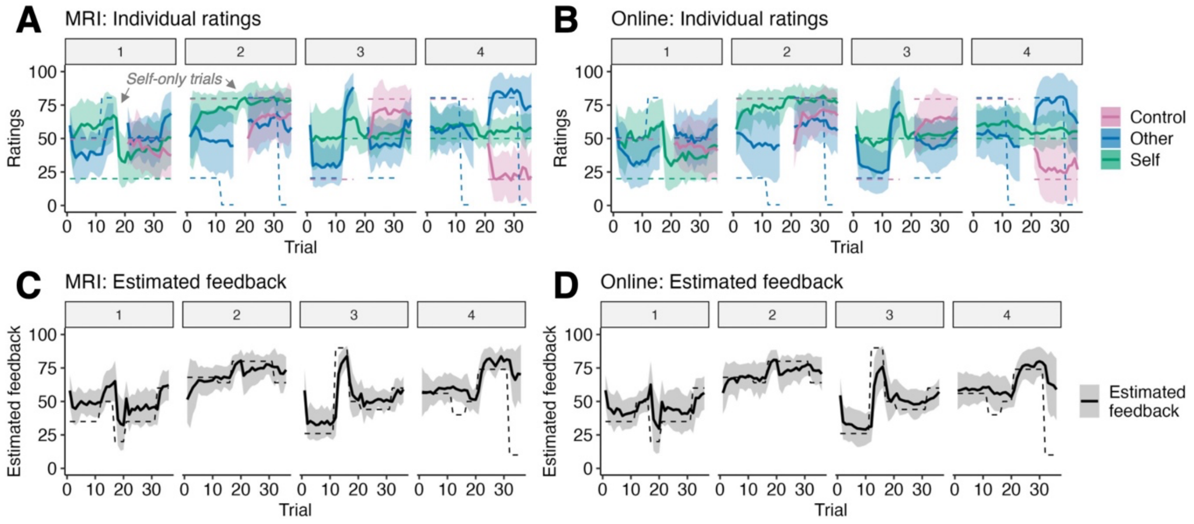


**Figure S3 (relating to Figure 2B-C).** Participants' average ratings and estimated feedback in each of the four task blocks. Shaded intervals are SD. Individual ratings of the **A)** MRI sample and **B)** Online sample. During the Self-only trials, when the feedback reflects only the self and participants only rated their own performance, their self estimates show a sudden adaptation to the true self level. Estimated feedback of the **C)** MRI sample and **D)** Online sample.

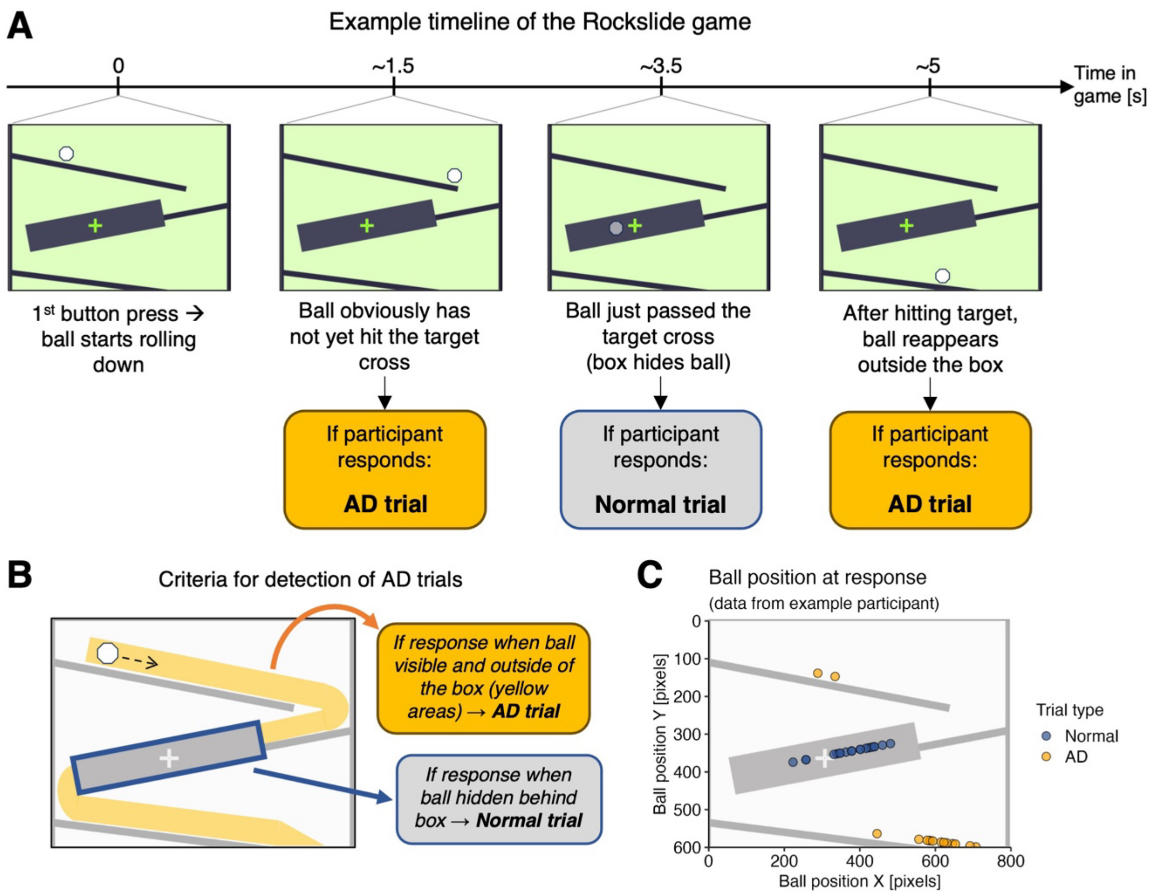


**Figure S4 (relating to Figure 3)**. Detection of AD trials when participants play the games. **A)** Example timeline in the Rockslide game. The game starts when participants press the button, which releases a ball that rolls down a set of slides. Depending on when participants respond (by button press), the trial is detected as normal or AD. Trials are detected as AD trials if participants make 'obvious' mistakes in the games. **B)** Specifically, this is the case if participants respond when the ball is still visible and outside of the box that hides the target cross. The trial is detected as a normal trial if participants respond when the ball is close to the target cross and hidden behind the box. **C)** Data from an example participant shows the clear difference between ball positions in AD and normal trials. Data is overlaid with an example screenshot of the rockslide game. Note that the blue dots (ball position on normal trials) vary in their positions because between trials, the target cross changed its position on the 2nd slide and participants could not see the ball behind the black box (and therefore were sometimes too early or too late in pressing the button).

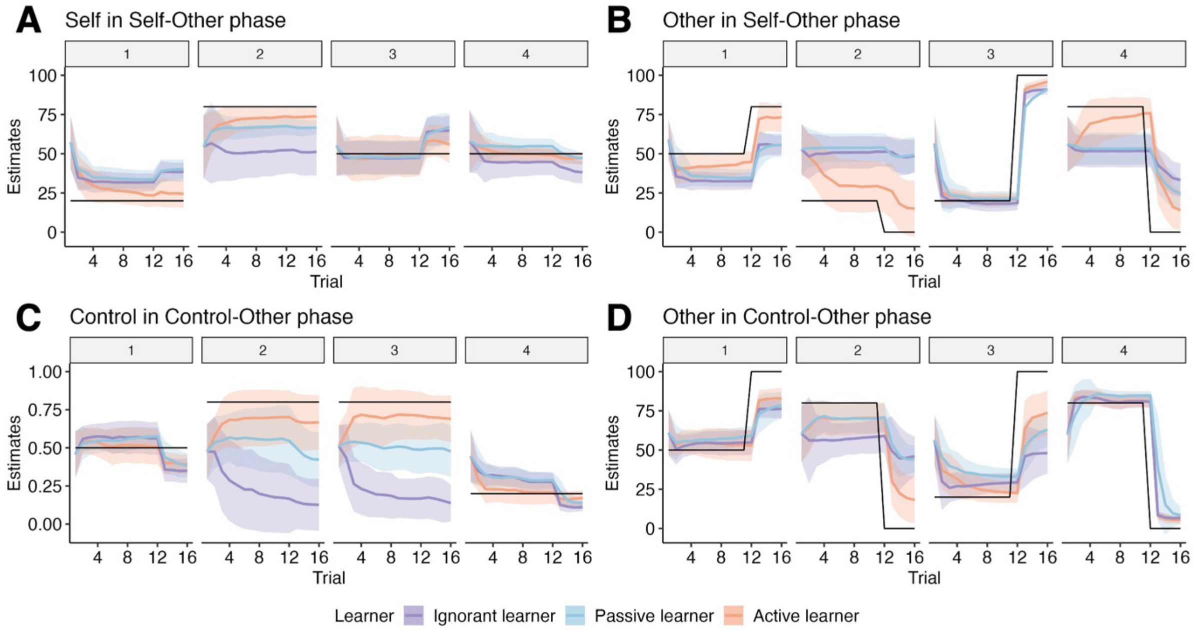


**Figure S5 (relating to Figure 3).** Average estimates per game block for the active, ignorant and passive Bayesian learner. Simulated data from MRI and online sample are pooled here. Lines are averaged estimates across participants, and shaded intervals are the standard deviation. Black lines indicate the true levels of each variable. **A)** Self estimates per game block in the Self-Other phase. **B)** Other estimates in the Self-Other phase. **C)** Control estimates in the Control-Other phase. **D)** Other estimates in the Control-Other phase. The active learner has lower rating errors than the suboptimal passive and ignorant learners (Main effect of learning model, $F(2,130)=447.24$, $p<0.001$, $\eta^2=0.83$; active vs. passive learner, $F(1,65)=461.54$, $p<0.001$, $\eta^2=0.72$; active vs. ignorant learner, $F(1,65)=485.83$, $p<0.001$, $\eta^2=0.86$; passive vs. ignorant learner, $F(1,65)=275.22$, $p<0.001$, $\eta^2=0.60$). Additionally, the active learner also has the lowest uncertainty at the end of the phases compared to the other suboptimal learning models (Main effect of learning model, $F(2,130)=325.07$, $p<0.001$, $\eta^2=0.79$; active vs. passive learner, $F(1,65)=350.98$, $p<0.001$, $\eta^2=0.79$; active vs. ignorant learner, $F(1,65)=351.19$, $p<0.001$, $\eta^2=0.72$; passive vs. ignorant learner, $F(1,65)=94.04$, $p<0.001$, $\eta^2=0.38$). Crucially, these results suggest that an optimal learner, given the knowledge of AD trials and the ability to learn from it, performs better (in terms of errors and uncertainty) than the two suboptimal learners, which either discard AD trials for learning (passive learner) or assume that AD trials are normal trials (ignorant learner). Summed rating errors and average uncertainty were extracted from the last phase trial of each phase in each block, and averaged across the individual ratings.

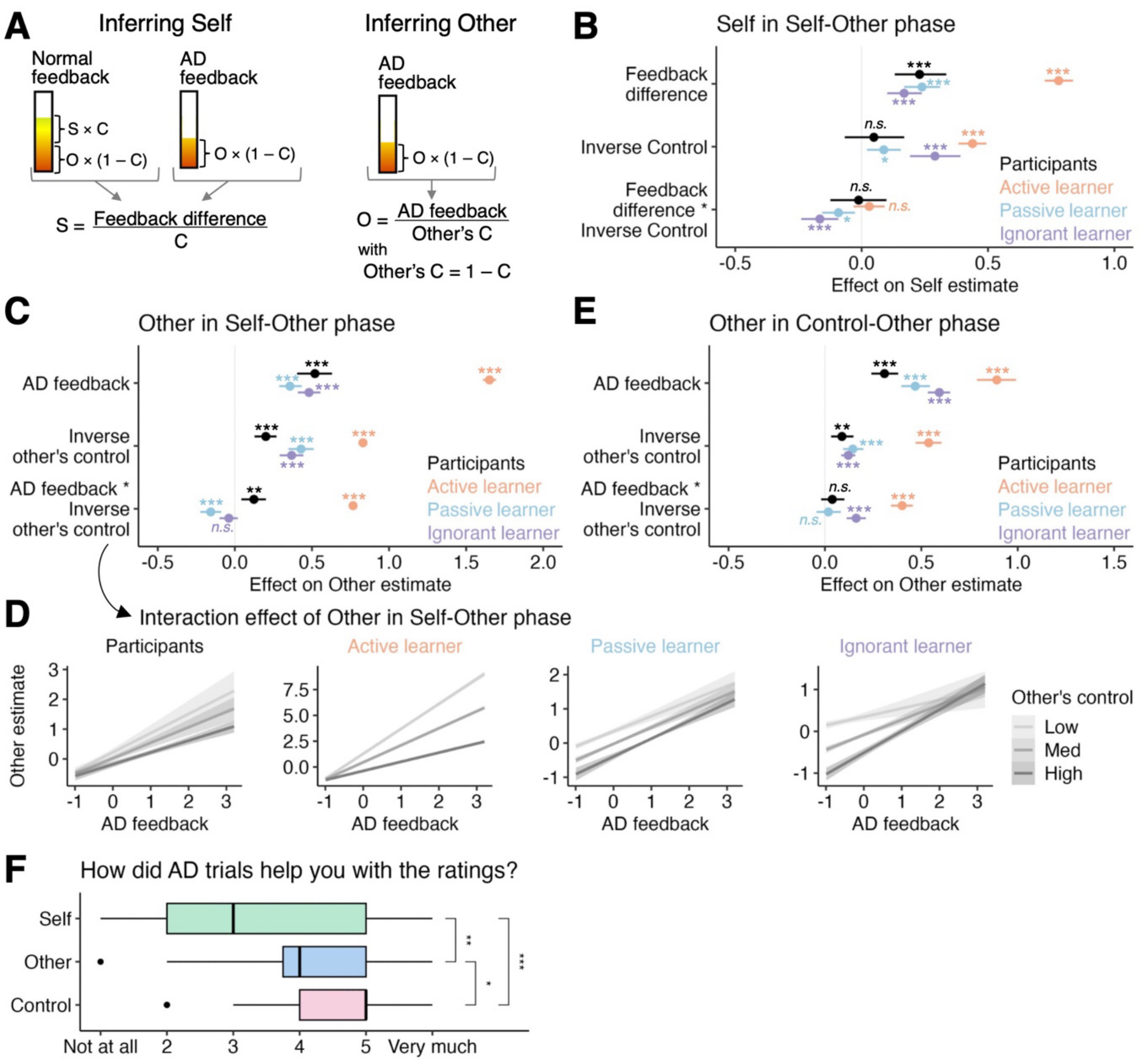


**Figure S6 (relating to Figure 4)**. Hierarchical regressions predicting how to assign credit from switch and AD trials to self and others. **A)** Optimally, in the Self-Other phase, self can be inferred from the feedback difference between normal and AD trials accounting for one's control. Note that this strategy is symmetrical to how control is inferred in the Control-Other phase (main text Figure 4). In both Self-Other and Control-Other phase, the other's performance can be inferred by just observing the AD feedback and accounting for the other's control (1-control). Note that in the Self-Other phase, the control is known while in the Control-Other phase, this requires the control which needs to be inferred as well. **B)** For inferring self-performance in the Self-Other phase, we found that participant performance neither resembled the active Bayesian learner (main effect of inverse control), nor the passive or ignorant Bayesian learning models (inverse control and interaction effect). This means that participants were not able to optimally extract the information afforded by AD and normal trials to infer their own performance. This is also reflected in their helpfulness ratings in the debrief questionnaire (main Figure 4C), where they reported AD trials as the least helpful for the self ratings. **C)** In contrast, participants inferred the other's performance similarly to the active Bayesian learner in the Self-Other phase. Here, the interaction effect AD feedback * Inverse other's control differentiated between, on one hand, participants and the active learner, and on the other hand, the two suboptimal learning models. Note however that participants' beta weights are not as close to the active learner as they are for the respective control inference (Figure 4D). Participants also rated AD trials as more helpful for learning about their control than learning about the other. For these reasons, we focussed our analyses on the control inference process and did not follow up this effect in inferring the other. **D)** The interaction effect from C shows that both participants and the Bayesian learner inferred that the other had a low performance if the AD feedback is low, irrespective of the other's control. If the AD feedback is high, however, they inferred a higher other performance level and even more so if the other had a low control because then the other's performance must account for the high AD feedback. The passive and ignorant Bayesian learners show markedly different behaviour in this interaction effect. Note that for easier interpretation, here, we relabelled the legend of "inverse other's control"

regressor to “Other’s control” and flipped its levels accordingly (e.g. “Low other’s control” is the effect of “high inverse other’s control”).**E)** In the Control-Other phase, we found that participants were not able to infer the other’s performance like an optimal active Bayesian learner (see interaction effect). **F)** In the debrief questionnaire, participants reported that they found the AD trials most helpful for inferring their controllability (ANOVA main effect of rating type (self, other, or control): $F(2,70)=13.57$, $p<0.001$, $\eta^2=0.17$; paired two-sided t-tests: self vs. other, $t(35)=-3.29$, $p=0.002$, $d=0.55$; self vs. control, $t(35)=-4.50$, $p<0.001$, $d=0.75$; other vs. control, $t(35)=-2.22$, $p=0.03$, $d=0.37$). Only data from the online sample is shown here because the MRI sample did not receive this debrief question. n=31 MRI, n=36 online; panels B-E: n.s., 95% CI includes 0; *, 95% CI excludes 0; **, 99% CI excludes 0; ***, 99.9% CI excludes 0; panel F: *, p<0.05; **, p<0.01; ***, p<0.001.

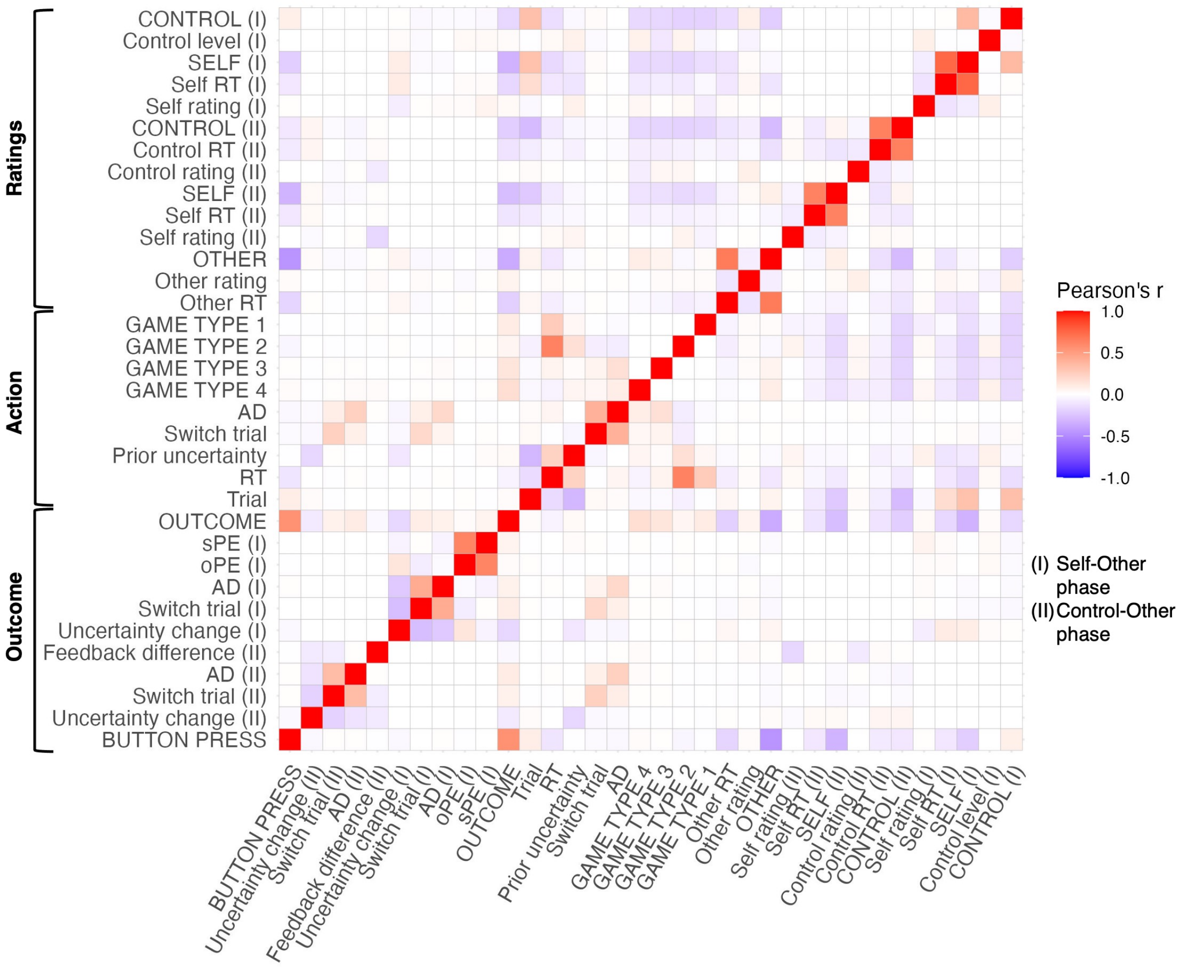


**Figure S7 (relating to Figure 5).** Correlations of regressors (convolved with hemodynamic response function) included in the fMRI whole brain analysis. Pearson's correlations were averaged across participants. Constant regressors are highlighted in uppercase. In the action phase when participants play the games, we included four constant regressors, one per game type (1=Light, 2=Slingshot, 3=Rockslide, 4=Bouncing ball).

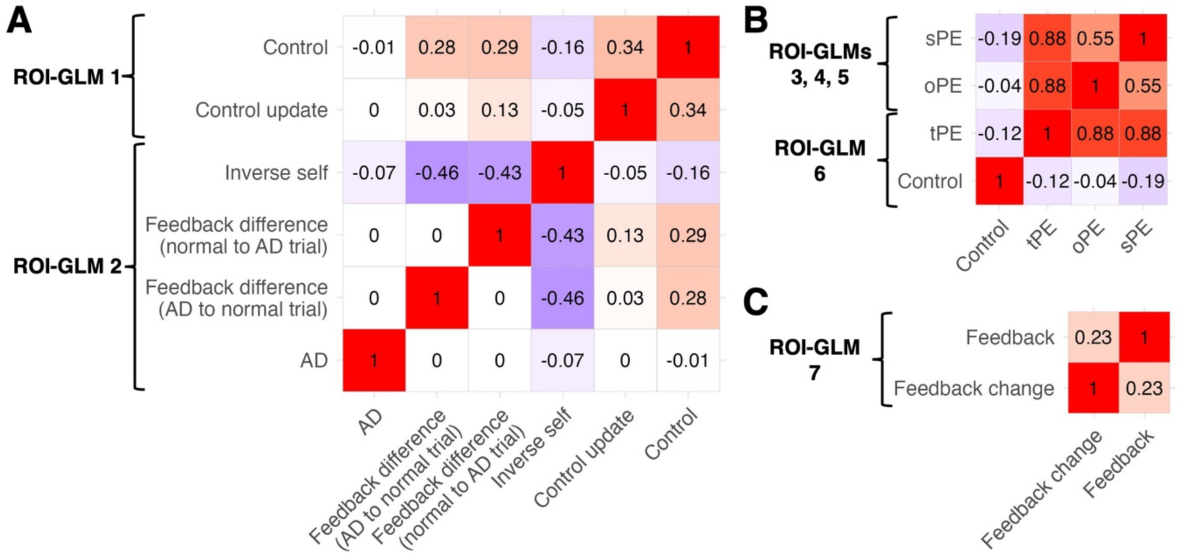


**Figure S8 (relating to Figures 6-7, S12).** Correlation matrices for regressors included in time course analyses (ROI-GLMs). **A)** Correlation of regressors shown in main Figure 6. Regressors for the ROI-GLMs 1 and 2 are shown in one correlation matrix to highlight that the regressors of control belief (ROI-GLM 1) and feedback differences (ROI-GLM 2) show low correlations (r<0.3). **B)** Correlations of regressors shown in main Figure 7. Since sPE and oPE showed a correlation of r=0.55, we tested the time courses of these regressors first separately (ROI-GLMs 3 and 4), and then together (ROI-GLM5). We also tested the effects of tPE and control separately in ROI-GLM6. **C)** Regressor correlations for supplementary Figure S12 (ROI-GLM7).

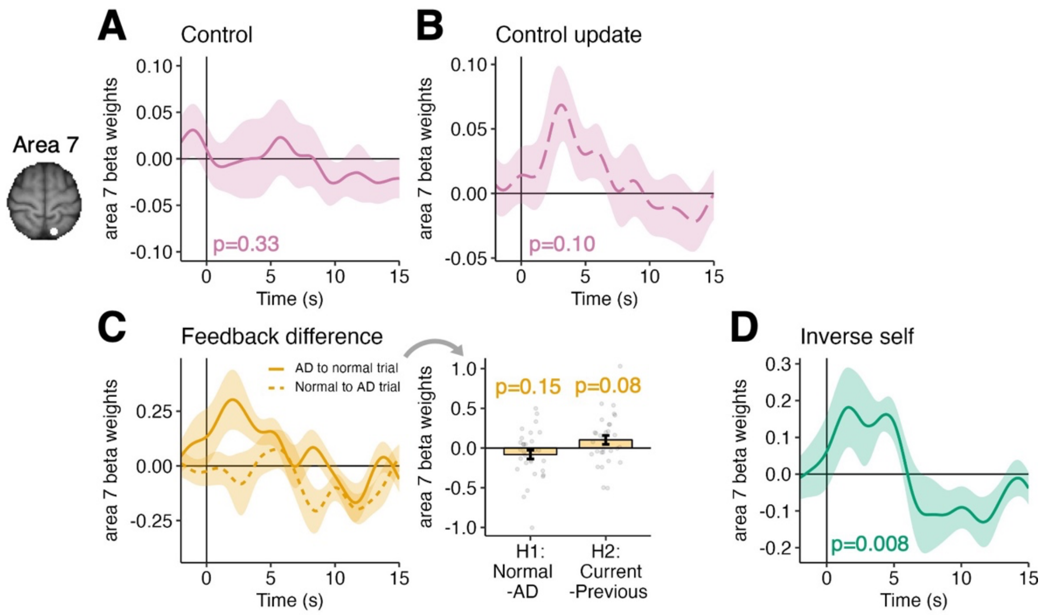


**Figure S9 (relating to Figure 6).** Time courses of feedback difference, control and self in left area 7, time-locked to the outcome phase. **A-B)** We found no evidence that area 7 activity signals the inferred control or the control update at the time of outcome (control, t(30)=-0.98, p=0.33; control update, t(30)=1.69, p=0.10; ROI-GLM1). **C)** While the waveforms of feedback differences look more alike H2 than H1, we found no evidence in favour of either hypothesis in area 7 (H1, t(30)=-1.48, p=0.15; H2, t(30)=1.83, p=0.08; ROI-GLM2). This is unlike the SMG result that we found (Figure 6E). **D)** In the same GLM, we also tested for the inverse self. We indeed found that area 7 tracks the prior inverse self estimate at the time of outcome (inverse self, t(30)=2.82, p=0.008; ROI-GLM2). n=31 MRI, mean beta weights are plotted as lines, with s.e.m. as shaded intervals.

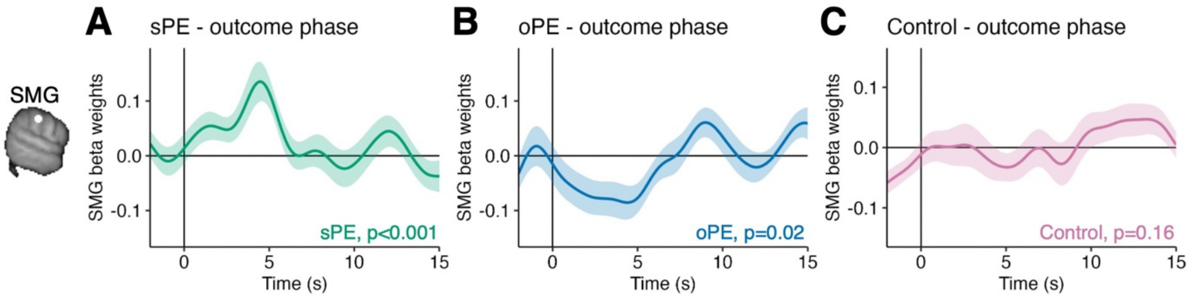


**Figure S10 (relating to Figure 7).** Time course analyses of sPE, oPE and Control in the SMG. **A)** Due to the correlation of sPE and oPE, we also ran a separate time course analysis including both sPE and oPE in one ROI-GLM (ROI-GLM5). Here, we found a significant effect of sPE (t(30)=3.83, p<0.001). **B)** oPE, which was included in the same GLM, also still showed a significant effect (t(30)=-2.41, p=0.02). **C)** In ROI-GLM6, we tested for effects of tPE (plotted in main Figure 7C) and control in the outcome phase. We did not find that SMG represents the known control level (t(30)=1.43, p=0.16) in this phase of the task – the Self-Other phase – when participants had already been instructed about its level and did not need to infer it. n=31 MRI, mean beta weights are plotted as lines, with s.e.m. as shaded intervals.

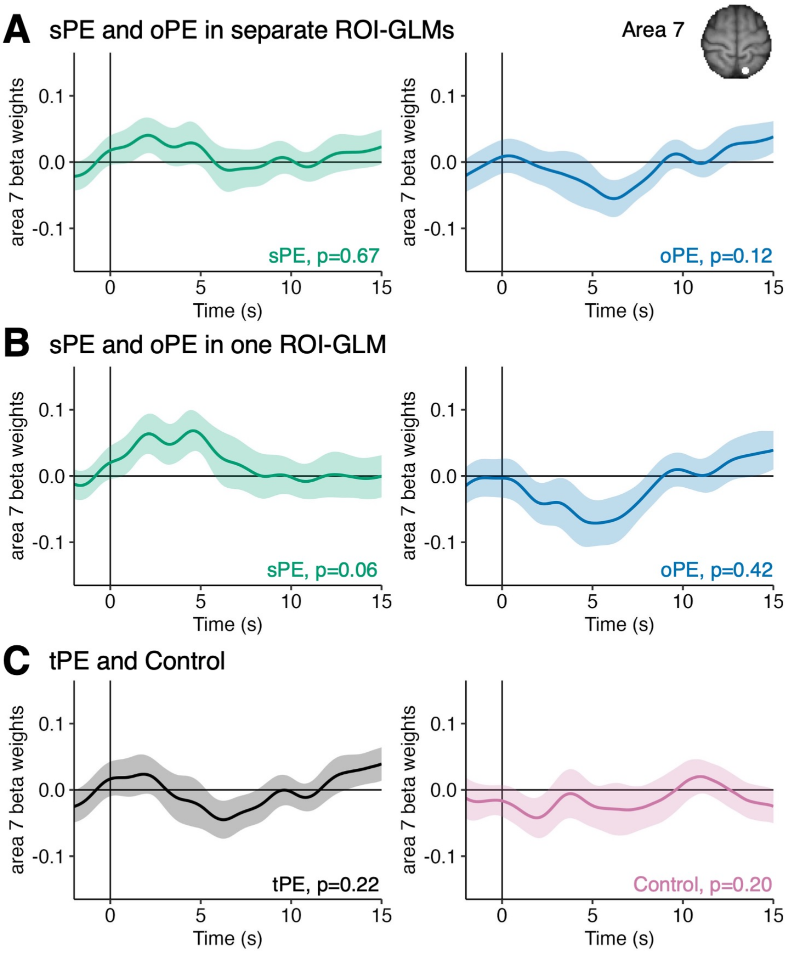


**Figure S11 (relating to Figure 7).** Time courses of sPE, oPE, tPE and control in area 7 at the time of outcome, during the Self-Other phase. **A)** We did not find that area 7 BOLD activity is significantly modulated by sPE nor oPE when tested in separate GLMs (sPE, $t(30)=0.43$, $p=0.67$, ROI-GLM3; oPE, $t(30)=-1.58$, $p=0.12$, ROI-GLM4). **B)** Tested in one GLM, sPE and oPE remained not significant in area 7 (sPE, $t(30)=1.92$, $p=0.06$; oPE, $t(30)=-0.81$, $p=0.42$; ROI-GLM5). **C)** BOLD activity in area 7 was not significantly modulated by tPE nor the known control either (tPE, $t(30)=-1.26$, $p=0.22$; control, $t(30)=1.30$, $p=0.20$; ROI-GLM6). n=31 MRI, mean beta weights are plotted as lines, with s.e.m. as shaded intervals.

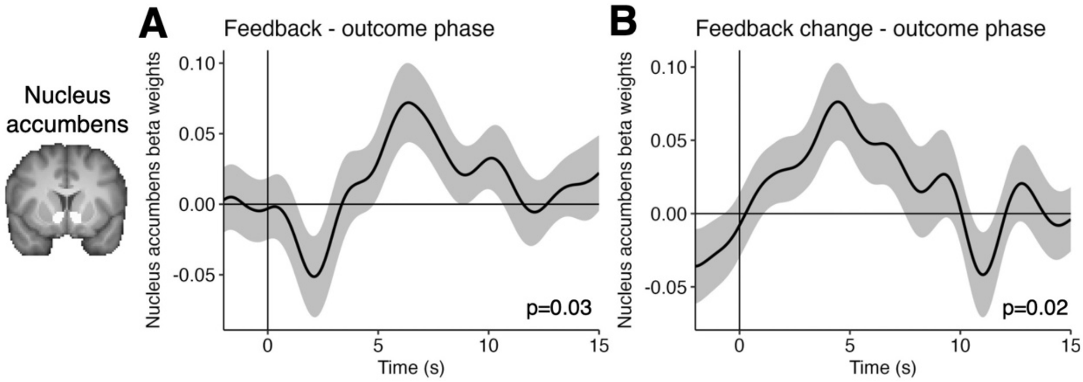


**Figure S12.** Time courses of feedback and feedback change in the ventral striatum in the outcome phase. Previous studies have shown that the ventral striatum is sensitive to reward (e.g. [3]). To test whether the ventral striatum also signals reward in our task, we ran a time course analysis on an anatomically defined bilateral mask of the nucleus accumbens (ROI-GLM7). **A)** We found that activity in the nucleus accumbens tracks the feedback that is being observed at the time of outcome ($t(30)=2.23$, $p=0.03$, ROI-GLM7). **B)** In the same GLM, we also found that the nucleus accumbens signals the change of the feedback (difference between current and last trial's feedback) at the time of outcome ($t(30)=2.48$, $p=0.02$, ROI-GLM7). Analyses were run on normal trials that followed normal trials, across Self-Other and Control-Other phase. The regressors show low correlations ($r=0.23$, see supplementary Figure S8C). n=31 MRI, mean beta weights are plotted as lines, with s.e.m. as shaded intervals.

#### Supplementary tables

Table S1 (relating to Figure 5). Peak coordinates of significant clusters in whole-brain fMRI analysis.

| Contrast | Region | Peak coordinates x/y/z (in mm MNI space) | z value |
|---|---|---|---|
| Uncertainty | Lateral occipital cortex, superior division | -38, -80, 32 | 4.7 |
| | Supramarginal gyrus (SMG) | -66, -24, 36 | 4.25 |
| | Lateral occipital cortex, inferior division | -50, -66, 2 | 4.14 |
| | Postcentral gyrus | -50, -38, 58 | 4.42 |
| | Area 7 | -12, -64, 66 | 3.94 |
| | Superior parietal lobule | -38, -46, 64 | 3.78 |
| | Intracalcarine Cortex | 2, -84, 2 | -4.77 |
| | Superior temporal gyrus, posterior division | -56, -34, 4 | -4.54 |
| | Lateral orbitofrontal cortex (lOFC) | -50, 22, 0 | -4.54 |
| | Occipital pole | 16, -90, 14 | -4.17 |
| | Dorsomedial prefrontal cortex (dmPFC) | -8, 52, 40 | -3.92 |
| | Temporal pole | -52, 12, -20 | -3.9 |
| Active disambiguation | Temporoparietal junction (TPJ)<br>- SMG subpeak (right): | 52, -44, 34<br><br>SMG: 64, -34, 30 | 5.82<br><br>SMG: 4.70 |
| | Dorsolateral prefrontal cortex (dlPFC)<br>- dmPFC subpeak: | Right: 28, 32, 34<br>Left: -44, 38, 28<br><br>dmPFC: 0, 40, 34 | Right: 4.93<br>Left: 3.76<br><br>dmPFC: 4.50 |
| | Precuneus | 2, -50, 46 | 4.98 |
| | SMG (left) | -64, -36, 32 | 5.04 |
| | Lateral frontal pole | 40, 52, 0 | 4.44 |
| | Inferior frontal gyrus, pars opercularis, stretching into lOFC (left) | 52, 18, -2 | 4.93 |
| | Lateral occipital cortex, inferior division | Right: 40, -86, -8<br>Left: -48, -74, 4 | Right: 4.08<br>Left: 3.98 |
| | Lingual gyrus | -6, -78, -6 | 4.65 |
| | Cerebellum, uvula | -26, -78, -24 | 4.80 |
| | 2 clusters in right lOFC (area 47o) | Cluster 1: 32, 22, -8<br>Cluster 2: 48, 24, -8 | Cluster 1: 4.57<br>Cluster 2: 4.23 |
| | Parahippocampal gyrus | 32, -44, -6 | 4.82 |
| | Cerebellum, tonsil | -38, -44, -44 | 3.82 |
| | Middle temporal gyrus | -64, -46, -2 | 3.70 |
| | Anterior cingulate cortex | 10, 50, 12 | 4.13 |
| | Cerebellum, anterior lobe | -32, -60, -32 | 4.11 |
| | Area 4 | Left: -42, -14, 50<br>Right: 44, -8, 58 | Left: -4.61<br>Right: -4.52 |
| | Supplementary motor area | 0, -4, 60 | -5.16 |

Family-wise error cluster corrected, $z > 3.1$, $p < 0.05$
